# Evidence that conserved essential genes are enriched for pro-longevity factors

**DOI:** 10.1101/2022.03.18.484929

**Authors:** Naci Oz, Elena Vayndorf, Mitsuhiro Tsuchiya, Samantha McLean, Lesly Turcios-Hernandez, Jason N. Pitt, Ben W. Blue, Michael Muir, Michael G. Kiflezghi, Alexander Tyshkovskiy, Alexander Mendenhall, Matt Kaeberlein, Alaattin Kaya

**Author notes:** equal contribution (Naci Oz, Elena Vayndorf).

## Abstract

At the cellular level, many aspects of aging are conserved across species. This has been demonstrated by numerous studies in simple model organisms like *Saccharomyces cerevisiae, Caenorhabdits elegans*, and *Drosophila melanogaster*. Because most genetic screens examine loss of function mutations or decreased expression of genes through reverse genetics, essential genes have often been overlooked as potential modulators of the aging process. By taking the approach of increasing the expression level of a subset of conserved essential genes, we found that 25% of these genes resulted in increased replicative lifespan in *S. cerevisiae*. This is greater than the ∼3.5% of genes found to affect lifespan upon deletion, suggesting that activation of essential genes may have a relatively disproportionate effect on increasing lifespan. The results of our experiments demonstrate that essential gene overexpression is a rich, relatively unexplored means of increasing eukaryotic lifespan.

## Introduction

Numerous studies have demonstrated that the genetic modifiers and molecular pathways altering aging are highly conserved across different phylogenies [1]. This suggests a hypothesis that genes that are evolutionarily constrained and functionally conserved may be more likely to alter the trajectory of the aging process. Specifically, we hypothesize that essential genes have high potential to alter the rate of aging. We use the term essential genes to refer to genes required for organismal survival. Essential genes tend to be conserved across species, and have fewer deleterious single-nucleotide polymorphisms [2]. Essential genes also tend to be highly expressed, often form stable complexes, and are relatively enriched for protein-protein interactions [2-4].

Conserved essential genes are relatively understudied in experimental aging research. By definition, null function for any essential gene results in inviability and, experimentally, reduced function is often associated with severe developmental deficits. Since most large-scale genetic studies of aging tend to utilize gene deletion or knock-down libraries (e.g., RNAi), essential genes have often been neglected. The effect of decreasing essential gene function on lifespan has been studied in *C. elegans* via partial knockdown by RNAi beginning in adulthood [5, 9]. These studies found that out of ∼2700 essential genes screened, reduced adult expression of only 2% extended lifespan [6, 7].

*Saccharomyces cerevisiae* has served as an important experimental organism for revealing essential gene functions, laying the foundation for understanding their functions in metazoans [10]. Perhaps due to their relatively compact genome, one in every six genes is reported to be essential for viability in *S. cerevisiae* (total 1,144 essential genes) [11, 12]. These conserved genes are so conserved they can often be swapped with orthologs from humans. Recent studies showed that 621 of the human homologs of yeast essential genes with diverse molecular functions can be functionally replaced by their human counterparts [13-16]. This finding indicates that the roles of conserved essential genes in the aging process may also be conserved across species.

To date, only a few essential genes have been reported to modulate replicative lifespan (RLS) in yeast. For example, the essential gene *LAG1* was found to increase RLS when overexpressed, and increased expression of its human homolog also increased RLS, despite low sequence similarity to the yeast protein [17]. Decreased activity of the guanine nucleotide exchange factor *CDC25* enhances lifespan in both yeast [18] and mice [19]. In addition, modulation of a conserved N-myristoyl transferase *NMT1* was shown to increase lifespan in yeast [20]. No large-scale analyses of essential gene effects on lifespan have been carried out in yeast due to the almost exclusive use of haploid gene deletion libraries, which do not include essential gene deletions (due to inviability).

A previous genome-wide scale comparative analysis of the 1805 known longevity-associated genes across 205 species revealed that longevity gene orthologs are substantially overrepresented, for the genes known to be essential for development and growth [21]. Based on these observations, we hypothesized that conserved essential genes may represent a largely untapped source of prolongevity genes. Specifically, given that reduced expression of only 2% of essential genes in *C. elegans* increased lifespan [6], we hypothesized that *increased expression* of essential genes may be more likely to increase lifespan. To test this idea, we initiated a systematic determination of the effect of increased essential gene dosage on RLS. Here we report the RLS results for 92 unique yeast strains each overexpressing a single conserved essential gene from a low copy plasmids. Our results suggest our hypothesis is correct; 23 (25%) of these strains showed significantly increased RLS. This frequency of RLS-extension appears to be much greater than the ∼4% of long-lived non-essential single-gene deletions. These essential prolongevity genes function in a variety of pathways and complexes. Our results demonstrate that increased expression of conserved essential genes is a relatively fruitful, and relatively unexplored, approach for extending lifespan.

## Results

### mRNA and protein levels of conserved essential genes are uncoupled during aging in *S. cerevisiae*

We used the WORMHOLE ortholog prediction tool [22] to identify *S. cerevisiae* orthologs for each of 1144 annotated yeast essential genes. Of these, 871 have orthologs in both nematode worms and humans **(Fig. 1, Table S1)**.

**Figure 1:**
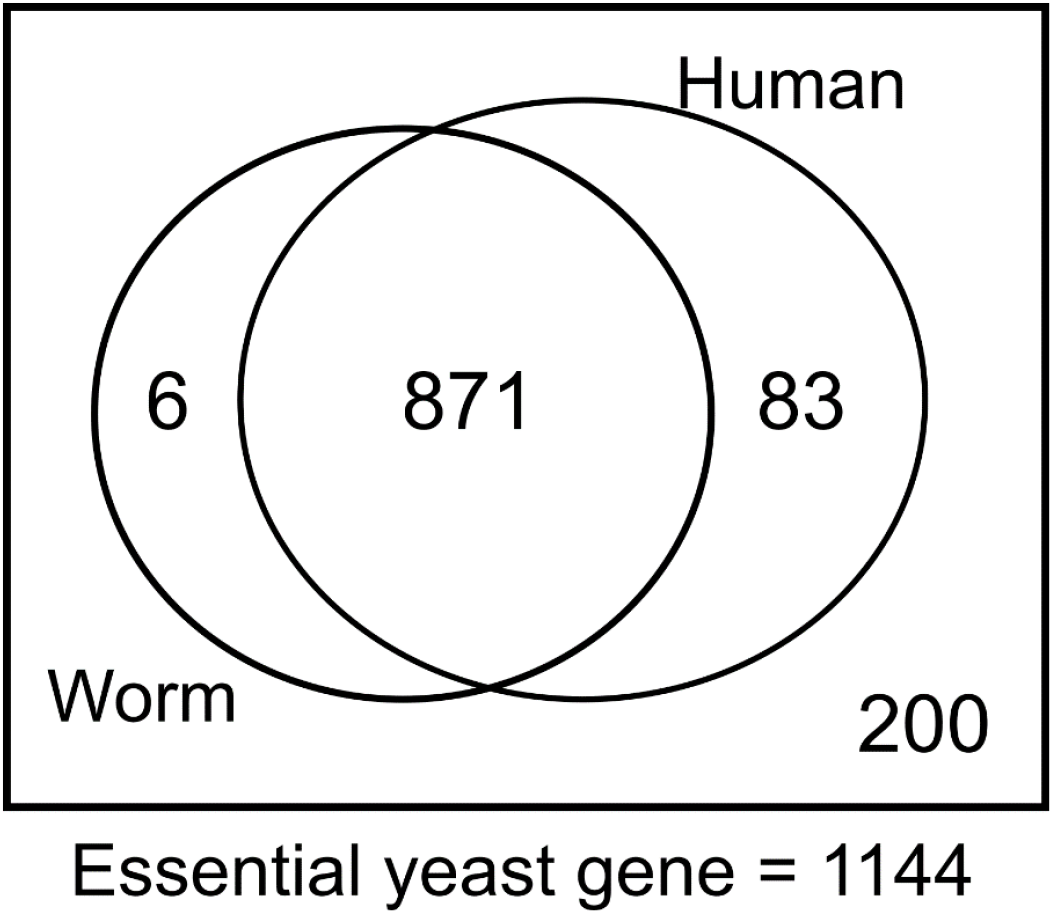
Number of yeast essential gene orthologs both in human and worm. Out of the 1144 essential genes analyzed, 871 of them have been found to have orthologs both in human and worm. It includes yeast essential genes with 1 to 1, 1 to many, many to many orthologs and homologs. Remaining 200 yeast essential genes have no homology in either of the species.

Next, to understand how essential genes are affected by aging at both transcript and protein levels, we analyzed age-dependent changes of their abundances. We obtained published data from yeast, in which age-dependent (replicative age) changes of 980 essential gene transcripts and 480 essential proteins were analyzed [23]. Our analysis revealed two main age-dependent clusters both for mRNA and protein abundances **(Fig. 2A)**. Overall, we identified 510 essential gene transcripts and 145 essential proteins with significant (Benjamini–Hochberg adjusted p-value ≤ 0.05) age-related changes (increased and decreased). Furthermore, we found no significant overlap between the expression patterns of proteins and their corresponding mRNAs (fisher exact test, p > 0.05) **(Fig. 2B)**. These data indicate an uncoupling between the mRNA and protein abundances of essential genes during aging and are consistent with a recent report suggesting that essential hub proteins likely play an important role in uncoupling of mRNA and protein abundances during replicative aging in yeast [24].

**Figure 2:**
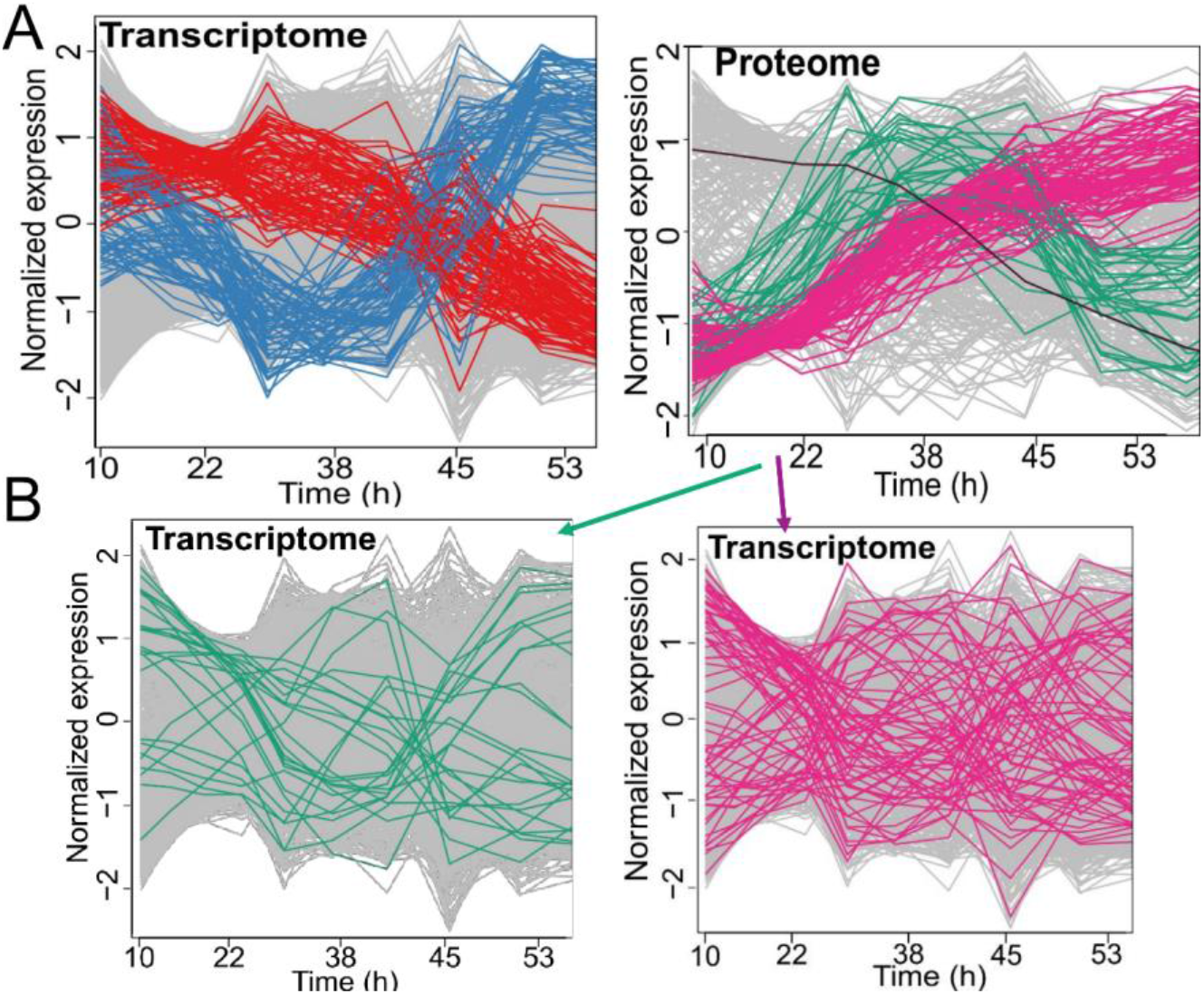
Age-dependent changes of essential gene abundances in yeast at mRNA and protein level. **(A)** Upper panels, shows two major clusters for age-dependent changes of essential genes at mRNA (left) and protein (right) level. Single black line at the protein panel is an example of one those 42 proteins that were found to have significant age-dependent decline. **(B)** Lower left panel shows the mRNA expression patterns of proteins (upper right) from the green cluster and the lower right panel shows the mRNA expression patterns of proteins from the pink cluster, suggesting uncoupling level between the mRNA and protein abundances of essential genes.

### Increased expression of essential genes increases RLS

In order to study the effect of essential gene overexpression on RLS, we utilized the recently developed unique Molecular Barcoded Yeast Open Reading Frame (MoBY-ORF-1) expression library [25, 26]. The MoBY-ORF 1 library was created by cloning each gene, controlled by its native promoter and terminator, into a low copy centromeric (*CEN*) vector. This vector carries a URA3 selectable marker along with two unique 20 bp long oligonucleotide barcodes (UPTAG and DNTAG) that can be amplified with universal primers, enabling cells carrying a specific ORF to be quantitatively detected by a single PCR reaction. The p5472-*CEN* plasmid persists in yeast cells with 1–3 copies per haploid genome and a centromeric sequence on this plasmid enables stable plasmid maintenance and correct plasmid distribution during cell division [27]. Overall, the MoBY-ORF1 library contains 993 individual glycerol stocks of bacterial strains, carrying uniquely barcoded essential ORFs on low copy yeast expression plasmids, representing ∼ 90% of the essential ORFs, as annotated in the *Saccharomyces* Genome Database [28]. We constructed a collection of 993 yeast strains each overexpressing a single essential gene by purification of plasmids from the individual MoBY-ORF1 library *E. coli* strains and individual transformation into the *S. cerevisiae* wild type BY4741 (WT) background (see materials and methods). An empty vector plasmid was transformed into BY4741 and serves as the control strain for all RLS experiments.

We selected a subset of 92 (**Table S1)** essential gene overexpressing strains for initial RLS analysis. After genotype verification of each MOBY-ORF1 clone, we first analyzed the effect of increased gene dosage on growth. Our analysis revealed that many of the strains had similar growth to the WT control cells harboring empty-vector (**Fig. 3A**). Only five strains showed severely affected growth from low copy essential gene overexpression, corresponding to the genes *PRP28, MED17, UTP4, RAD3* and *BFR2* (**Fig. 3A)**.

**Figure 3:**
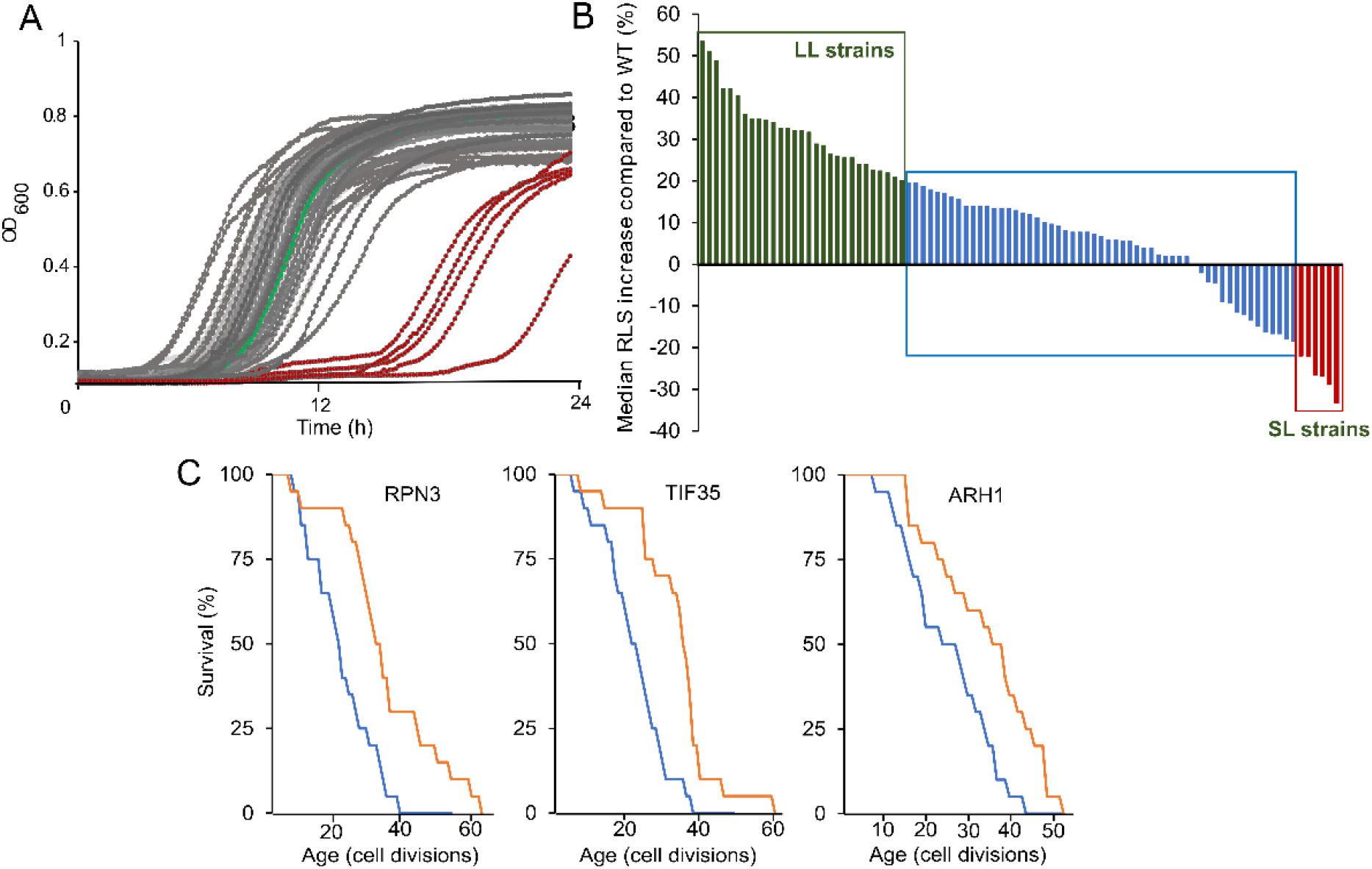
Growth and median RLS change across the strains tested in the pilot RLS analysis. **(A)** Green curve represents the WT control and gray curves represent the strains harboring essential gene expression constructs. The five strains with decreased growth rate (*PRP28, MED17, UTP4, RAD3* and *BFR2)* are indicated with red curves. **(B)** Median RLS changes in comparison to the WT control cell were plotted for 92 strains. Green lines represent long lifespan strains (LL) that increased the median RLS by at least 20%, as blue bars indicate ambiguous strains with no significant median RLS changes. Red bars represent short lifespan strains (SL) that decreased the median RLS by at least 20%. **(C)** RLS analysis for three LL strains after plasmid loss. Blue lines indicate mortality curves for the strains after losing plasmids and orange lines indicate mortality curves with increased gene dosage through overexpression with plasmid, which extend RLS significantly (p ≤ 0.05).

Initial RLS analysis revealed that ∼32% of the strains (30/92) showed a greater than 20% increase in median RLS while ∼7% of the strains (6/92) showed greater than 20% decrease in median RLS, compared to matched control cells (**Fig. 3B, Table S1)**. Statistical analysis of observed lifespan differences (Wilcoxon Rank Sum Test) revealed that 23 out of 92 strains (25%) significantly extended median RLS (p ≤ 0.05) in comparison to the WT control (**Table S1)**. To further confirm that the observed longevity phenotype is directly related to increased expression of the essential gene, we treated three of the long-living strains with 5-FOA to induce plasmid loss (see materials and methods). After verification of plasmid loss in these strains (*RPN3, ARH1, TIF35*), we repeated the RLS experiment and found that lifespan for each of these strains was reverted back to the wild type (WT) control, confirming the increased essential gene dosage was the cause of the observed longevity phenotype **(Fig. 3C)**.

### RNA-seq of five essential gene overexpression strains reveals both distinct and overlapping transcriptional signatures of longevity

To gain insight into the signatures of essential-gene mediated lifespan extension at the transcript level, we performed RNA-Seq analysis on five of the identified long-lived strains **(Fig. 4A)**. These selected strains overexpressed: *LSM5* (part of Lsm1p complex involved in mRNA decay, human homolog *LSM5*), *SNU13* (part of U3 snoRNP involved in rRNA processing, human homolog *SNU13*), *TIF35* (subunit of the core complex of eIF3, human homolog *EIF3G*), *UTP6* (component of the processome, containing the U3 snoRNA that is involved in processing of pre-18S rRNA, human homolog *UTP6*) or *PRP22* (DEAH-box RNA-dependent ATPase/ATP-dependent RNA helicase, human homolog *DHX8*), respectively. First, in order to determine how much each strain increased expression level, we quantified the increase in expression for each essential gene, based on their mRNA level. We found that overexpression of these genes using their native promoter and a low copy plasmid caused 1.97 to 3.85-fold increase in mRNA levels. **(Fig. 4B)**.

**Figure 4.**
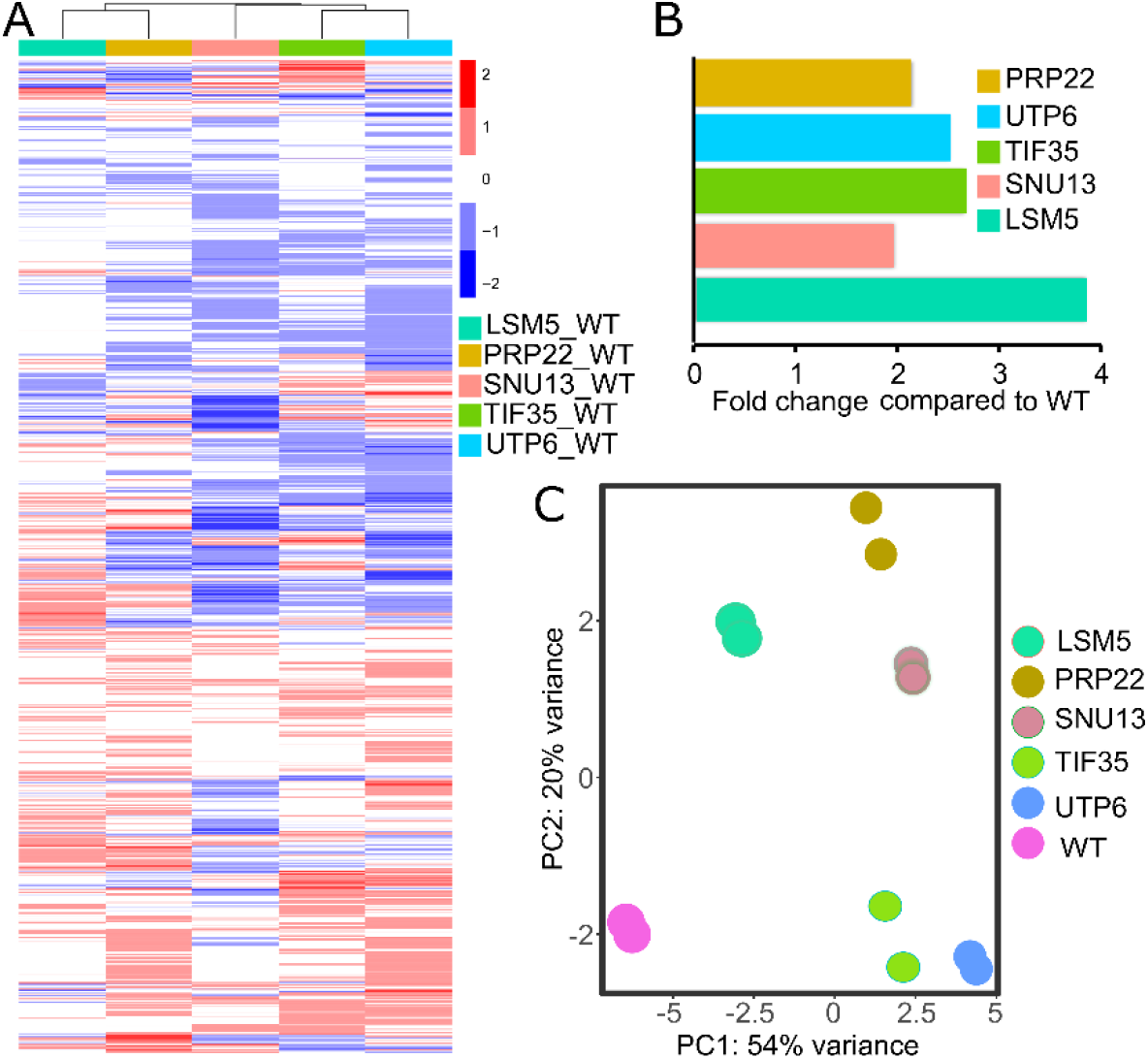
Gene expression profiles across the subset of long-lived strains: **(A)** Heat map on the left panel shows genome-wide transcriptional changes of 5807 genes (log2 fold) in strains with increased dosage of *LSM5, PRP22, TIF35, SNU13* and *UTP6*.compared to the WT control. For each strain, an average of two biological replicates represented. Red bins indicate increased and blue bins indicate decreased mRNA expression for the genes. Spearman correlation matrix of aggregated gene expression profiles across strains was used for clustering (top panel on the heatmap). **(B)** mRNA level of corresponding genes overexpressed under their own promoter and terminators. **(C)** PCA representing variances across the strains based on the transcriptional changes.

Next, we analyzed genome-wide transcriptional changes of these strains by comparing them to the WT and generated expression heat maps **(Fig. 4A)**. After filtering and quality control, our dataset contained RNA-seq reads for 5,807 transcripts across the five long-lived strains **(Table S2)**. Overexpression of *LSM5* caused the lowest number of transcript alterations, significantly changing 126 transcripts. *UTP6* overexpression caused the highest number of transcript alterations, changing 274 transcripts, compared to the WT control (adj. p ≤ 0.05; fold change >=1.5). Overexpression of *UTP6, TIF35* and *SNU13* induced statistically significant overlap (Spearman correlation test, adj. p value = 2.9×10^−7^), their transcriptional changes associated with expression patterns shared more than 34% of up- and 43% of downregulated genes, respectively **(Table S2)**.

To further understand similarities in gene expression variation across these strains, we performed a principal component analysis (PCA), based on normalized read counts for each strain. We found that overexpression of *UTP6, TIF35* and *SNU13* clustered together, suggesting similar transcriptional changes **(Fig. 4C)**. Furthermore, these three strains shared 74 genes that significantly changed expression (fold change > = 1.5, adj. p ≤ 0.05) **(Table S2)**. Functional enrichment analysis revealed a significantly enriched set of 21 downregulated and five upregulated proteins involved in RNA-mediated transposition (adj. p = 6.8×10^−16^).

The data suggest a hypothesis that increased expression of *UTP6, SNU13* and *TIF35* might promote longevity by decreasing the activity of mobile DNA elements (transposable elements), which have been linked to the aging process in various models [29, 30]. Functional enrichment analysis of the strain overexpressing *LSM5* revealed significant (adj. p ≤ 0.05); upregulation of carbohydrate metabolic process and cellular glucose homeostasis and down regulation of transcription by RNA polymerase II. Significant enrichment (adj. p ≤ 0.05) for upregulation of spliceosome and mRNA surveillance pathway and down regulation of citrate cycle and pyruvate metabolisms were observed in *PRP22* overexpressing strain.

Overall, our RNA-seq data indicates even a 2-fold increase in mRNA level of essential genes is sufficient to initiate changes in cellular processes by rewiring genome-wide transcriptional patterns. Such changes are a rich, untapped source for understanding genetic mechanisms of longevity determination.

## Discussion

This study addresses a question that has been relatively unstudied in experimental aging research: What is the role of conserved essential genes in the aging process? From an initial analysis of 92 strains each overexpressing a single essential gene in yeast, we observed a surprisingly high frequency of lifespan extension in 23 cases. The genes we identified have diverse functions. These data suggest that conserved essential genes may play a heretofore unappreciated central role in longevity determination. If orthologs of these genes similarly regulate longevity in multicellular eukaryotes, this will have a profound impact on the field from both a theoretical and practical perspective, potentially creating a new paradigm for how we think about conserved genetic control of longevity, in terms of the roles of essential genes.

Although systematic gene knockdown of essential genes has been studied in *C. elegans* [5, 6], there have been no large-scale efforts to explore the effects of essential gene overexpression in the context of aging biology in any organism. The prior emphasis on decreased gene function [8, 9, 31-37] has likely led to a bias in the field toward studying genes and pathways whose normal function promotes aging and limits lifespan.

The increased RLS observed in 25% of the yeast strains examined here is striking in comparison to prior studies of gene knockdown/deletion in both yeast and worms. In the previous RNAi screen of ∼2700 essential worm genes, lifespan extension was observed ∼2.4% (64 genes) of the time [6]. In a genome-wide screen of non-essential gene deletions for yeast RLS, lifespan extension was observed ∼3.4% of the time. Thus, conserved essential gene overexpression may increase lifespan much more frequently than gene knockdown/deletion.

It remains to be determined whether overexpression of orthologs of the genes identified here will increase lifespan in other organisms. Prior work has indicated that genetic control of longevity is significantly conserved between yeast and worms [7, 10], but these studies have been restricted to RNAi/deletion analyses. It will be of considerable interest to determine whether overexpression of orthologs to the genes identified here can increase lifespan in *C. elegans*. Given that we have only analyzed less than 10% of the conserved essential yeast genes and have already identified more than 20 novel longevity factors in yeast, we predict that a comprehensive analysis of conserved essential gene overexpression in yeast followed by ortholog testing in worms could yield several novel conserved longevity factors, most if which also have mammalian orthologs.

Among the long-lived strains we identified are two proteasome related genes, four genes involved in mRNA surveillance, five vesicle trafficking and secretory pathway genes, two post translation modification genes, and three genes involved in DNA repair. There are also individual genes which play a role in diverse cellular processes such as oxidative stress resistance, chromatin structure remodeling complex, and iron-sulfur (Fe-S) cluster biogenesis **(Table S1)**.

Our analysis revealed several exceptionally long-lived strains whose RLS exceeded most of the long-lived gene deletions identified from more than 5000 strains analyzed [31]. In particular two genes, *GDI1* and *TIF35* both extended median RLS by more than 50% **(Fig. 5)**. *GDI1* (human ortholog: *GDI2*) is a GDP dissociation inhibitor involved in ER-Golgi vesicle-mediated transport [38, 39]. Another longevity gene, *TIF35* is a subunit of the core complex of eukaryotic translation initiation factor 3 (eIF3) involved in ribosomal scanning during translation reinitiation [40]. This is among the largest effect sizes ever reported for year RLS, comparable to the very long-lived *Δhxk2 Δfob1* double mutant and *Δhxk2 Δsir2 Δfob1* triple mutant cells, known to act on both the genetically-distinct SIR2/rDNA and CR pathways [1]. ER-Golgi vesicle-mediated transport plays a role in pre mRNA/protein quality control and autophagy, a recycling mechanism that has been associated with lifespan regulation in different organisms [39, 42-44]. Identification of several genes from these pathways (*TRS31, TRS35 SEC20, SEC26, RPN3, PRE1, PUP3*) in our RLS screen further supports the role autophagy and proteasome function in lifespan regulation. In addition, expression of four small nucleolar RNA (snoRNA) genes (*UTP6, UTP7, SNU13, SYF1*), also increased median RLS up to 34% and maximum RLS up to 69% **(Table S1)**. These essential genes are involved in chemical modifications of other RNAs, mainly ribosomal RNAs, transfer RNAs and other small nuclear RNAs [45, 46]. This data suggests that mRNA translational fidelity and post-transcriptional mRNA modification machinery, a process which also directly influences the <u>proteostasis</u> capacity of a cell might be also associated with increased lifespan in those long living strains as it was suggested previously [47, 48]. Overall, these data highlight known, and unknown potential new regulatory pathways associated with the function of essential genes in the regulation of aging.

**Figure 5:**
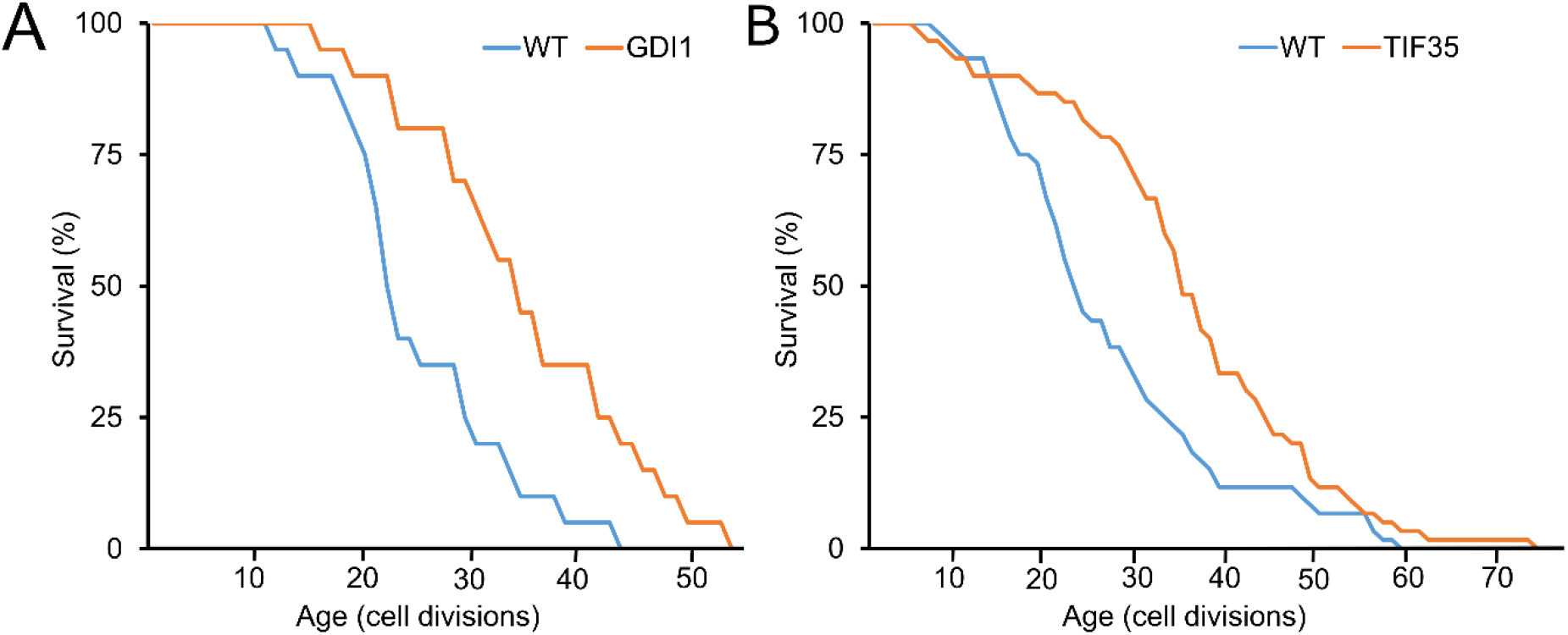
The increased dosage effect of GDI1 and TIF35 on RLS: Increased gene dosage of **(A)** GDI1 and **(B)** TIF35 significantly (p ≤ 0.05) extends median RLS. Red line represents overexpression lines and the gray line represents WT control harboring an empty vector backbone. Number of mother cells analyzed for each group along with p values can be found in Table S1.

Essential genes tend to have a higher mRNA level and encode highly abundant proteins that engage in more protein*-*protein interactions [2-4, 49]. Consistent with our data, it has been shown that abundance of most of the essential genes do not show significant age-dependent changes at both the transcript and the protein level [24]. However, gradual decay or loss of interactions to essential proteins in the protein-protein interaction networks as a function of aging in yeast has been previously reported [24]. It should also be noted that many of the known longevity genes are involved in the protein-protein interaction complexes, in which hubs tend to correspond to essential genes [3, 24, 50, 51]. Essential proteins are known to have long half-lives [52]. It is possible that essential long-lived proteins may maintain their abundances in old cells even though their translation levels decline. However, these essential genes probably have age-associated functional impairments because of accumulated damages, (e.g., protein oxidation, carbonylation, translational errors) [53]. Our approach of mild increases of their abundances via overexpression might replace those damaged essential proteins with functional ones and restore those protein-protein interaction complexes late in life. Such stochastic changes in transcript level of essential genes late in life might regulate transcript and protein level of those interacting partners and associated pathways as well.

Overall, we believe that this study is the first ever focused on studying the systematic effects of essential genes on lifespan in budding yeast. Because overexpression of essential genes has never been studied before in yeast aging, every hit represents a newly discovered longevity gene with a conserved human ortholog. In theory, classification of the candidate lifespan regulators may represent new molecular mechanisms underlying the lifespan variation with perturbation of conserved essential gene functions, which might also lead to therapeutic targets. Further studies will yield a wealth of new information about the genetic and molecular determination of aging in yeast and will identify dozens of candidate genes for testing in high eukaryotes.

## Materials and Methods

### Yeast strains and culture conditions

We utilized the MoBY-ORF1 expression library [25, 26] to express our essential gene of interest in laboratory BY4741 WT cells. This library is provided in bacterial stock; therefore we first prepared plasmids from these bacterial strains and transformed them into BY4741 WT cells. The MoBY-ORF1 library already provides us individual constructs, each harboring one of the essential genes of interest controlled by its native promoter and terminator, into a low copy centromeric (*CEN*) vector. We sequenced all MoBY-ORF1 constructs, tested in RLS analysis, to verify insertion of correct clones and six of the incorrect gene constructs re-cloned from the BY4741 genome into the same vector backbone using the Gibson assembly (NEB E5510S), a similar method by which the MoBY-ORF1 library was created [54]. Individual colonies from each yeast transformation were selected and kept as a frozen glycerol stock that all the RLS experiments and the molecular phenotyping of long living strains was performed from. WT cells, harboring an empty vector backbone (p5472CEN) of MOBY1-ORF1 library, was used as a control for RLS analysis. For testing the growth effect of increased dosage of each essential gene, strains were cultured overnight in a 96-well plate incubator at 30 °C in-URA medium. Next day, 1 μl from overnight culture was transferred to the YPD and growth was monitored in 96-well plate using Epoch2 (BioTek, Winooski, VT, USA) kinetic growth analyzer by analyzing optical density of OD_600_.

### Yeast lifespan analysis

RLS was determined using a modification of our previously published protocol [55]. Yeast cell cultures for each strain were freshly started from frozen stocks on – URA plates and grown for 2 days at 30 °C prior to dissections. Several colonies were streaked onto new YPD with 2 % glucose. For each strain, 20 to 80 individual mother cells were analyzed in RLS analysis. In each experimental group of RLS analysis, the same number of mother cells from WT control (BY4741-p5472CEN) were also analyzed. RLS of each strain was determined by comparing the result to their corresponding WT control from the same experimental group. All the RLS experiments were performed in a blind fashion, with strains assigned with random numbers to eliminate any potential bias during the RLS assay. To test the plasmid dependence of an observed lifespan phenotype, the URA3 marker was used for quick elimination of the plasmid from the host cell by treating them with 5-FOA [56].

### RNA-seq analysis

Two independent cultures for each yeast strain harboring overexpression cassette of *LSM5, TIF35, UTP6, PRP22* or *SNU13* along with WT control and three independent cultures for young and old cell populations of strain harboring overexpression cassette of ARH1 and WT control were collected at the OD_600_ = 0.4 on -URA medium to isolate RNA from each culture using Quick-RNA 96 Kit from Zymo Research (Cat. number: R1053). To prepare RNA-seq libraries, Illumina TruSeq RNA library preparation kits were used according to the user manual, and RNA-seq libraries were loaded on Illumina HiSeq 4000 platform to produce 150 bp paired-end sequences. After quality control and adapter removal, the HISAT2 software package [57] was used to map the reads against the S288c reference genome. Read counts per gene were calculated using featureCounts [58]. To filter out genes with low numbers of reads, we used filterByExpr function from the edgeR package. Differential expression analysis was performed using the DESeq2 R package [59]. The resulting P values were adjusted using the Benjamini and Hochberg’s approach for controlling the false discovery rate.

## Acknowledgement/Funding

This work was supported by NIH/NIA 1K01AG060040 to A.K. Studies performed by M.K. were funded by a University of Washington Nathan Shock Center of Excellence in the Basic Biology pilot award to A. K. (P30AG013280). The Project was also supported by a Longevity Impetus Grant from Norn Group to MK. The funders had no role in study design, data collection and interpretation, or the decision to submit the work for publication.

## Data Availability

Transcriptome (RNA-sequencing) reads and fragments per kilobase of transcript per million mapped read values deposited in the Gene Expression Omnibus (GEO) database, www.ncbi.nlm.nih.gov/geo (accession no. XXXX).

